# Metabolite profiling and cytotoxic activity of Andean potatoes: polyamines and glycoalkaloids as potential anticancer agents in human neuroblastoma cells *in vitro*

**DOI:** 10.1101/2022.10.13.512083

**Authors:** María Luciana Lanteri, María Ximena Silveyra, Mónica Mariela Morán, Stéphanie Boutet, Deyvis-Dante Solis-Gozar, François Perreau, Adriana Balbina Andreu

**Author notes:** Corresponding author E-mail address (M.L. Lanteri).

## Abstract

Andean potatoes (*Solanum tuberosum* L. ssp. *andigena*) are a good source of dietary antioxidant polyphenols. We have previously demonstrated that polyphenol extracts from Andean potato tubers exerted a dose-dependent cytotoxic effect in human neuroblastoma SH-SY5Y cells, being skin extracts more potent than flesh ones. In order to gain insight into the bioactivities of potato phenolics, we investigated the composition and the *in vitro* cytotoxic activity of total extracts and fractions of skin and flesh tubers of three Andean potato cultivars (Santa María, Waicha, and Moradita). Potato total extracts were subjected to liquid-liquid fractionation using ethyl acetate solvent in organic and aqueous fractions. We analyzed both fractions by HPLC-DAD, HPLC-ESI-MS/MS, and HPLC-HRMS. Results corroborated the expected composition of each fraction. Organic fractions were rich in hydroxycinnamic acids (principally chlorogenic acid isomers), whereas aqueous fractions contained mainly polyamines conjugated with phenolic acids, glycoalkaloids, and flavonoids. Organic fractions were not cytotoxic against SH-SY5Y cells, and indeed, some increased cellular metabolism compared to controls. Aqueous fractions were cytotoxic and even more potent than their respective total extracts. Treatment with a combination of both fractions showed a similar cytotoxic response to the corresponding extract. According to correlation studies, it is tempting to speculate that polyamines and glycoalkaloids are crucial in inducing cell death. Our findings indicate that the activity of Andean potato extracts is a combination of various compounds and contribute to the revalorization of potato as a functional food.

## 1. Introduction

Potato (*Solanum tuberosum*) is the main non-cereal food crop worldwide. It is the most significant contributor to daily dietary antioxidants (Chun et al., 2005), and considered a functional food. Potato is classified into two subspecies, *andigena* and *tuberosum*, the former characterized by a great diversity of tuber forms and colors (Hijmans & Spooner, 2001).

Potatoes are rich in many phytochemicals derived from the specialized metabolism with diverse structures and biological functions. Phenylpropanoids are a group of thousands of compounds, including hydroxycinnamic acids (HCCs) (Ezekiel et al., 2013). Flavonoids are divided into six main subclasses: flavanols, flavonols, flavones, isoflavones, flavanones, and anthocyanins (Lewis et al., 1998). Polyamines occur primarily as putrescines, spermines, and spermidines, and can be conjugated with phenolic acids (Huang et al., 2017). Glycoalkaloids are plant steroids, being chaconine and solanine the main ones found in potatoes (Friedman, 2006).

In plants, phytochemicals participate in diverse processes like protection against biotic and abiotic stresses. They also exhibit diverse health-promoting effects such as antiinflammatory, anticarcinogenic, proapoptotic, and antioxidant activities (Friedman, 1997). Taking into account the growing interest in studying natural bioactive compounds, potato tubers and their industrial byproducts appear as promising sources suitable for application in the food and pharmaceutical industries. Metabolites can induce antitumoral effects based on the type, concentration, and how they interplay. Some studies have determined that polyphenol extracts from fruits and vegetables, including potatoes, provide antiproliferative activity against multiple cancer cell lines with null or minimum effect against normal cells (Bontempo et al., 2013; Vizzotto et al., 2014; Ombra et al., 2015; Kubow et al., 2017). Specifically, extracts of potato tubers inhibited proliferation in the prostate (Reddivari et al., 2007; Bontempo et al., 2013), stomach (Hayashi et al., 2006), liver (Wang et al., 2011), and colon (Madiwale et al., 2011; Kubow et al., 2017) cancer cells. Further, potato extracts rich in anthocyanins (Bontempo et al., 2013) or glycoalkaloids (Friedman et al., 2005; Nogawa et al., 2019) showed a dose-dependent antitumor activity in cervical, breast, and leukemia cancer cells.

Previously, we studied the cytotoxic effect of Andean potato polyphenol extracts against human neuroblastoma and hepatocarcinoma cell lines. Treatments resulted in a dose-dependent viability reduction in all cell types (Martínez et al., 2018; Silveyra et al., 2018). However, we do not know which compound(s) exert(s) these biological activities. So far, few studies have addressed potato bioactivity and nutraceutical composition in fractions or semi-purified extracts (Sánchez Maldonado et al., 2014; Chaparro et al., 2018; Nogawa et al., 2019).

This work aimed to fraction and to characterize the metabolites in polyphenol extracts of skin and flesh Andean potato cultivars (Santa María, Waicha, and Moradita) and analyze their cytotoxic effects against human neuroblastoma SH-SY5Y cells. We applied liquid-liquid fractionation and mass spectrometry to identify the compounds responsible for the bioactivity.

## 2. Materials and methods

### 2.1. Plant material

Moradita, Waicha, and Santa María potato cultivars were selected based on previous work from our laboratory (Silveyra et al., 2018) and other groups (Calliope et al., 2018). The skin and flesh colors are purple and yellow for Moradita, pink and yellow for Waicha, and intensely red for Santa María. Potato cultivars were grown in a field located in Yavi Department (22°6’4”S, 65°35’44”O, 3377 MAMSL), Jujuy, Argentina, during the 2012/2013 campaign. All cultivars were planted in random plots and harvested at the end of their respective cycles. We pooled skin and flesh of freshly harvested tubers to generate a representative sample for each cultivar. The material was immediately frozen in liquid nitrogen and stored at −80°C until analysis. Frozen potato pieces were freeze-dried and finely powdered.

### 2.2. Extraction and fractionation of total extracts (TEs)

Lyophilized tissues (2 g DW) from potato tuber skin or flesh were incubated overnight with 40 mL 100% methanol at 4°C in darkness with constant agitation (Figure 1). The methanolic extracts were centrifuged at 6000g for 20 min at 4°C, and thoroughly dried in a rotary evaporator (Senco) followed by a vacuum concentrator (Martin Christ). We obtained ethanolic extracts by adding 2.6 mL ethanol 80% (v/v) and centrifugation at 13000g for 10 min. Liquid-liquid fractionation of the ethanolic extracts was done according to the protocol described in Oki et al. (2002). Briefly, we separated 0.8 mL as total extracts (TEs), and we added 40 μL of trifluoroacetic acid and 1.6 mL of distilled water to the remaining volume (1.8 mL). The ethanol was removed by evaporation in a rotary vacuum concentrator. Then, four consecutive extractions were done with 1.8 mL ethyl acetate each. The resulting organic fractions (OFs), aqueous fractions (AFs), and TEs were dried in a vacuum concentrator and dissolved in methanol 30% v/v (except for HPLC-ESI-MS/MS and HPLC-HRMS analysis) with proportional volumes according to the DW of each fraction (0.5 mL for TEs, and 1.125 mL for OFs and AFs) (Figure 1).

**Figure 1.**
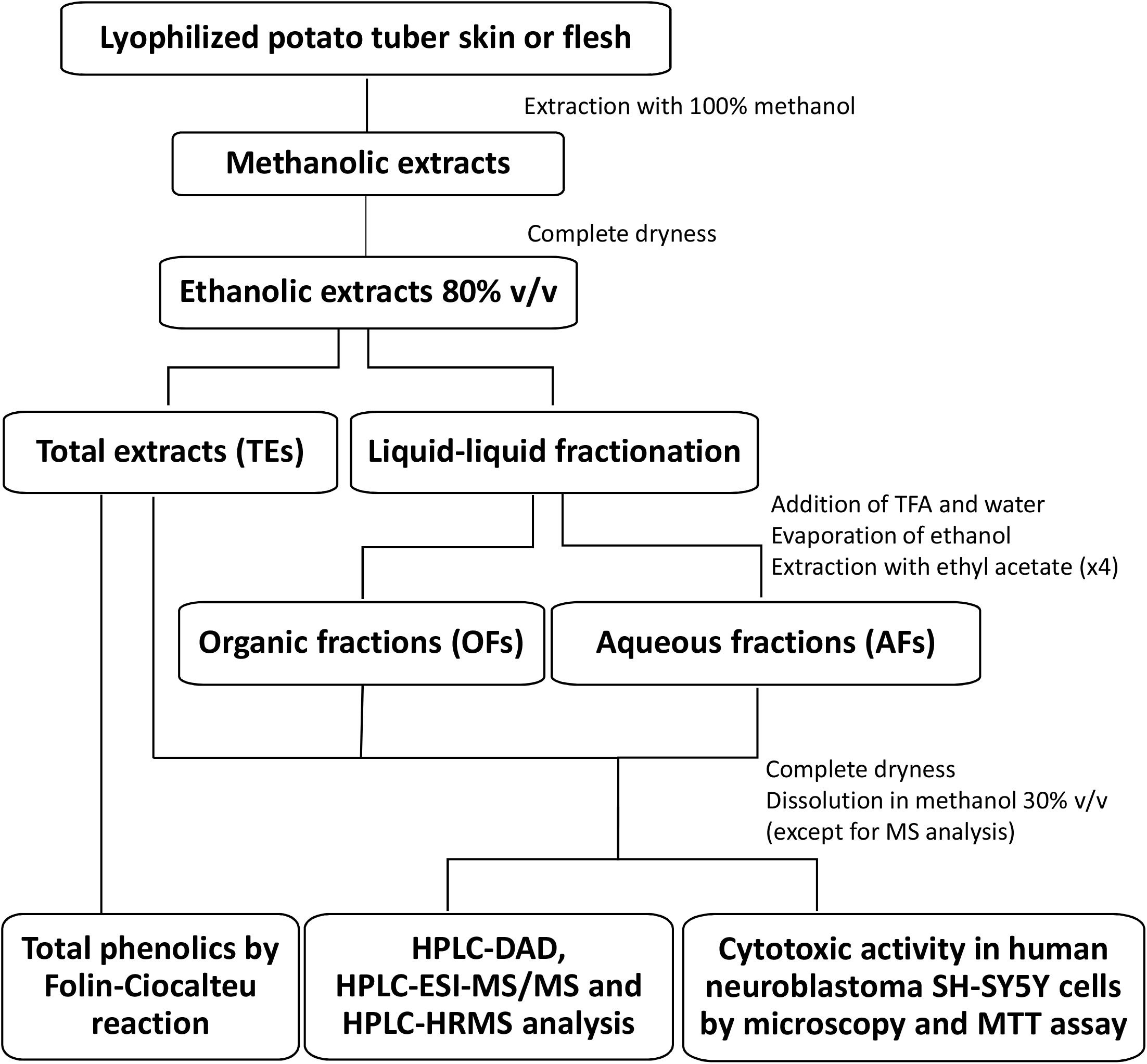
Simplified scheme for the obtention of potato extracts, the protocol of fractionation, and other assays done. TFA, trifluoroacetic acid.

### 2.3. Analysis by HPLC-DAD

Quantification of HCCs and anthocyanidins, present in TEs and fractions of each cultivar and tissue, was carried out using a Shimadzu LC-Solution system equipped with a diode array detector (DAD) as described in Silveyra et al. (2018). All peaks and standards were integrated according to the baseline. We expressed the levels of metabolites as μg/g DW in three biological replicates.

### 2.4. Analysis by HPLC-ESI-MS/MS for the relative quantification and HPLC-HRMS for the annotation of compounds

The dry pellets of TEs, OFs, or AFs were dissolved in methanol/H_2_O/acetone/trifluoroacetic acid (40/32/28/0.05, v/v). Then, 50 ng of apigenin (Extrasynthese, Genay, France) was added as an internal standard, and extracts were filtered through Whatman TM filter paper grade GF/A. After this, extracts were introduced in the ESI source of a U-HPLC Acquity equipped with a PDA detector (Waters, Milford, USA). Separation was achieved on a reverse-phase column Uptisphere C18 ODB, 150x2.1 mm (Interchim, Motluçon, France) using a flow rate of 0.4 mL/min and a binary gradient: (A) acetic acid 0.1% (v/v) in water, and (B) acetic acid 0.1% (v/v) in acetonitrile. The gradient program was 0-2 min 95% A, 2-4 min 90% A, 4-17 min 40% A, 17-23 min 100% A, and 23-25 min 95% A. The analysis was performed on a Xevo TQS triple quadrupole mass spectrometer (Waters, Milford, USA) operating in different scanning modes to identify the compounds and full scan for the semi-quantification. Relevant parameters were set as follows: capillary 2.70 kV, extractor 3 V, source block, and desolvation gas temperatures 150°C and 600°C, respectively. Nitrogen was used to assist nebulization and desolvation (7 Bar and 1000 L/h, respectively), and argon was used as collision gas for the MS/MS mode at 2.83x10^-3^ mBar. Levels of metabolites are expressed in μg apigenin equivalents/g DW based on the extracted ion chromatogram of each compound. The same samples were analyzed by HPLC-HRMS to confirm the annotation using standards or the accurate mass.

### 2.5. Total phenolics

The total phenolic content of TEs was determined by the Folin-Ciocalteu method and expressed as μg 5-chlorogenic acid (5-CGA) equivalents/mL in three biological replicates, according to Martínez et al. (2018). Blank samples consisted of 30% methanol (v/v).

### 2.6. Cytotoxic measurements and microscopy

Human neuroblastoma SH-SY5Y cells (ATCC CRL-2266) were maintained in DMEM/F-12 (Sigma-Aldrich) as described in Silveyra et al. (2018). After 18 h of treatments with different fractions, cell cultures were photographed under an Olympus CKX41 inverted microscope equipped with an Olympus Q-Color 3 camera (Figure 1). The % cell viability was obtained using the MTT assay (Sigma-Aldrich) and normalizing to the values in the absence of treatments. We presented all data as mean ± SD. The extracts were tested in triplicate, and the experiment was performed at least three times.

### 2.7. Statistical analysis

We used GraphPad software (Prism) to perform the statistical analysis for cell viability assays using One way ANOVA multiple comparisons followed by Dunnet’s test. We considered significant differences at P values ≤0.05 between treatments and the control with 30% methanol (v/v). Pearson product-moment correlation coefficients (R^2^ values) were calculated using Microsoft Excel with the mean values of metabolites levels and % cell viability.

## 3. Results

### 3.1. Fractionation of TEs: recovery and purity of OFs and AFs

To estimate the fractions’ recovery and purity, we calculated the % of compounds (HCCs or anthocyanidins) related to the total amount obtained for each TE by HPLC-DAD analysis (Table 1). Results indicate that the fractionation of TEs was successful because the OFs were rich in HCCs, and the AFs contained the most of the anthocyanidins. The % recovery of HCCs in OF was ≥51.1% in all the fractions, whereas the % recovery of anthocyanidins in AF was ≥87.5%. In addition, no more than 18.6% of recovered HCCs remained in the AFs, whereas a maximum of 21.7% of the recovered anthocyanidins contaminated the OFs. In short, AFs were purer than OFs (Table 1).

**Table 1.** The total content of HCCs and anthocyanidins in TE, OF, AF, and OF+AF from tuber skin and flesh of Santa María, Waicha, and Moradita cultivars. Levels of metabolites were determined by HPLC-DAD and expressed as μg/g DW. Values of OF+AF are theoretical and correspond to the sum of OF and AF in each case. Percentages between parentheses represent the recovery to the corresponding TE. ND, not detected.

| Cultivar | Compounds | Skin |  |  |  | Flesh |  |  |  |
| --- | --- | --- | --- | --- | --- | --- | --- | --- | --- |
|  |  | TE | OF | AF | OF+AF | TE | OF | AF | OF+AF |
| Santa María | HCCs | 2453.61<br>(100) | 2018.67<br>(82.3) | 264.31<br>(10.8) | 2282.98<br>(93.0) | 1663.41<br>(100) | 1322.72<br>(79.5) | 34.50<br>(2.1) | 1357.22<br>(81.6) |
|  | Anthocyanidins | 581.95<br>(100) | 89.99<br>(14.9) | 544.55<br>(93.6) | 631.54<br>(108.5) | 345.37<br>(100) | 32.03<br>(9.3) | 390.41<br>(113) | 422.44<br>(122) |
| Waicha | HCCs | 666.28<br>(100) | 340.54<br>(51.1) | 96.12<br>(14.4) | 436.66<br>(65.5) | 373.98<br>(100) | 267.63<br>(71.6) | 69.50<br>(18.6) | 337.13<br>(90.1) |
|  | Anthocyanidins | 60.41<br>(100) | 13.09<br>(21.7) | 58.78<br>(97.3) | 71.87<br>(118.9) | ND | ND | ND | ND |
| Moradita | HCCs | 1809.54<br>(100) | 1609.11<br>(88.9) | 228.71<br>(12.6) | 1837.82<br>(101.6) | 104.43<br>(100) | 94.26<br>(90.3) | 7.32<br>(7.0) | 101.58<br>(97.3) |
|  | Anthocyanidins | 158.21<br>(100) | 11.87<br>(7.5) | 138.37<br>(87.5) | 150.24<br>(95) | ND | ND | ND | ND |

OF+AF values are theoretical and represent the sum of individual OF and AF. These values are a reference to know how close the values of the sum of the fractions are compared to the TE values. OF+AF values were between 65.5% and 122% of the corresponding TE (Table 1).

### 3.2. HPLC-DAD chromatograms and composition of TEs, OFs, and AFs

We quantified 12 individual metabolites (six HCCs and six anthocyanidins) in each cultivar, tissue, and fraction with HPLC-DAD analysis. The composition of each extract and its fractions is displayed in the chromatograms (Supplemental Figures 1-3); however, we could not identify some of the peaks. Profiles at 320 nm for HCCs in all TEs were similar to those obtained in OFs and differed from those obtained in AFs (Supplemental Figures 1-3). Besides, the anthocyanidin profiles at 510 nm were similar between TEs and AFs (Supplemental Figures 1-3). We expected this because HCCs were recovered in OFs, and anthocyanins were recovered in AFs. The HPLC-DAD chromatograms display different mAU scales because some samples were diluted before injection. The corrected values multiplied by the dilution factor are reported in Tables 1 and 2.

**Table 2.** Content of individual HCCs and anthocyanidins in TE, OF, AF, and OF+AF from skin and flesh of the three Andean potato cultivars. Levels of metabolites were determined by HPLC-DAD and expressed as μg/g DW. ND, not detected.

| Santa María |  |  |  |  |  |  |  |  |
| --- | --- | --- | --- | --- | --- | --- | --- | --- |
| Compounds | Skin |  |  |  | Flesh |  |  |  |
|  | TE | OF | AF | OF+AF | TE | OF | AF | OF+AF |
| 5-CGA | 2056.91 | 1656.57 | 100.08 | 1756.65 | 1495.93 | 1188.89 | 24.34 | 1213.24 |
| 4-CGA | 217.89 | 219.51 | 87.80 | 307.32 | 121.14 | 97.72 | 8.78 | 106.50 |
| 3-CGA | 121.14 | 112.20 | 76.42 | 188.62 | 46.34 | 36.10 | 1.38 | 37.48 |
| CA | 57.67 | 30.40 | ND | 30.40 | ND | ND | ND | ND |
| CouA | ND | ND | ND | ND | ND | ND | ND | ND |
| FA | ND | ND | ND | ND | ND | ND | ND | ND |
| **Total HCCs** | **2453.61** | **2018.67** | **264.31** | **2282.98** | **1663.41** | **1322.72** | **34.50** | **1357.22** |
| Delphinidin | ND | ND | ND | ND | ND | ND | ND | ND |
| Cyanidin | 2.28 | ND | 2.28 | 2.28 | 1.46 | ND | 1.14 | 1.14 |
| Petunidin | ND | ND | ND | ND | ND | ND | ND | ND |
| Pelargonidin | 468.29 | 73.33 | 429.27 | 502.60 | 241.63 | 32.03 | 262.11 | 294.15 |
| Peonidin | 111.38 | 13.66 | 113.01 | 126.67 | 94.47 | ND | 99.02 | 99.02 |
| Malvidin | ND | ND | ND | ND | 7.80 | ND | 28.13 | 28.13 |
| **Total anthocyanidins** | **581.95** | **86.99** | **544.55** | **631.54** | **345.37** | **32.03** | **390.41** | **422.44** |
| **Total** | **3035.56** | **2105.67** | **808.86** | **2914.53** | **2008.78** | **1354.75** | **424.91** | **1779.66** |

| Waicha |  |  |  |  |  |  |  |  |
| --- | --- | --- | --- | --- | --- | --- | --- | --- |
| Compounds | Skin |  |  |  | Flesh |  |  |  |
|  | TE | OF | AF | OF+AF | TE | OF | AF | OF+AF |
| 5-CGA | 217.28 | 106.71 | 15.65 | 122.36 | 190.41 | 155.69 | 25.20 | 180.89 |
| 4-CGA | 189.63 | 93.70 | 27.24 | 120.93 | 109.11 | 76.50 | 19.35 | 95.85 |
| 3-CGA | 176.73 | 72.15 | 48.78 | 120.93 | 70.73 | 33.09 | 24.63 | 57.72 |
| CA | 59.37 | 62.65 | 4.46 | 67.11 | 0.21 | 0.10 | 0.28 | 0.37 |
| CouA | 1.49 | 0.79 | ND | 0.79 | 0.61 | 0.75 | ND | 0.75 |
| FA | 21.79 | 4.54 | ND | 4.54 | 2.91 | 1.50 | 0.04 | 1.54 |
| **Total HCCs** | **666.28** | **340.54** | **96.12** | **436.66** | **373.98** | **267.63** | **69.50** | **337.13** |
| Delphinidin | ND | ND | ND | ND | ND | ND | ND | ND |
| Cyanidin | ND | ND | ND | ND | ND | ND | ND | ND |
| Petunidin | ND | ND | ND | ND | ND | ND | ND | ND |
| Pelargonidin | 34.80 | 8.46 | 33.50 | 41.95 | ND | ND | ND | ND |
| Peonidin | 9.51 | 1.30 | 8.94 | 10.24 | ND | ND | ND | ND |
| Malvidin | 16.10 | 3.33 | 16.34 | 19.67 | ND | ND | ND | ND |
| **Total anthocyanidins** | **60.41** | **13.09** | **58.78** | **71.87** | **ND** | **ND** | **ND** | **ND** |
| **Total** | **726.69** | **353.63** | **154.90** | **508.53** | **373.98** | **267.63** | **69.50** | **337.13** |

| Moradita |  |  |  |  |  |  |  |  |
| --- | --- | --- | --- | --- | --- | --- | --- | --- |
| Compounds | Skin |  |  |  | Flesh |  |  |  |
|  | TE | OF | AF | OF+AF | TE | OF | AF | OF+AF |
| 5-CGA | 1484.15 | 1322.36 | 128.86 | 1451.22 | 85.37 | 75.20 | 7.32 | 82.52 |
| 4-CGA | 171.14 | 144.31 | 59.76 | 204.07 | 18.70 | 18.29 | ND | 18.29 |
| 3-CGA | 65.04 | 52.03 | 39.43 | 91.46 | ND | ND | ND | ND |
| CA | 88.91 | 90.19 | ND | 90.19 | ND | 0.76 | ND | 0.76 |
| CouA | 0.31 | 0.23 | 0.66 | 0.89 | ND | ND | ND | ND |
| FA | ND | ND | ND | ND | 0.37 | ND | ND | ND |
| **Total HCCs** | **1809.54** | **1609.11** | **228.71** | **1837.82** | **104.43** | **94.26** | **7.32** | **101.58** |
| Delphinidin | 0.49 | ND | 0.49 | 0.49 | ND | ND | ND | ND |
| Cyanidin | 0.16 | ND | 0.49 | 0.49 | ND | ND | ND | ND |
| Petunidin | 142.76 | 9.11 | 122.28 | 131.38 | ND | ND | ND | ND |
| Pelargonidin | ND | 1.46 | 0.33 | 1.79 | ND | ND | ND | ND |
| Peonidin | 4.88 | 0.49 | 5.53 | 6.02 | ND | ND | ND | ND |
| Malvidin | 9.92 | 0.81 | 9.27 | 10.08 | ND | ND | ND | ND |
| **Total anthocyanidins** | **158.21** | **11.87** | **138.37** | **150.24** | **ND** | **ND** | **ND** | **ND** |
| **Total** | **1967.76** | **1620.98** | **367.08** | **1988.07** | **104.43** | **94.26** | **7.32** | **101.58** |

For each cultivar, skin extracts showed higher levels and more diversity of compounds than the corresponding flesh extracts (Table 2). Concerning HCCs in TEs, Santa María skin had the highest total concentration (2453.61 μg/g DW with 2056.91 μg/g DW 5-CGA), followed by Moradita skin (1809.54 μg/g DW with 1484.15 μg/g DW 5-CGA). Waicha skin had a total concentration of 666.28 μg/g DW, with 217.28 μg/g DW 5-CGA, 189.63 μg/g DW 4-CGA and 176.73 μg/g DW 3-CGA. Among flesh extracts, Santa María contained the highest total HCCs concentration (1663.41 μg/g DW), followed by Waicha and Moradita (373.98 μg/g DW and 104.43 μg/g DW, respectively). Again, all the HCCs were represented in Waicha flesh (Table 2).

Regarding the anthocyanidins, Santa María skin presented the highest abundance (581.95 μg/g DW total, 468.29 μg/g DW pelargonidin). Waicha skin total concentration of anthocyanidin was 60.41 μg/g DW, with 34.80 μg/g DW corresponding to pelargonidin. Moradita skin concentration was 158.21 μg/g DW, composed primarily of 142.76 μg/g DW petunidin. Finally, Santa María flesh total concentration was 345.37 μg/g DW, comprised principally of 241.63 μg/g DW pelargonidin. No anthocyanidins were found in Waicha nor Moradita flesh extracts (Table 2).

### 3.3. Tentative identification of metabolites by HPLC-ESI-MS/MS and HPLC-HRMS

In order to gain information about the nature of anthocyanins and the presence of other compounds, we performed untargeted metabolomics for potato extracts. Table 3 shows the HPLC-ESI-MS/MS data for 30 compounds organized by retention time (RT) and divided into six families. We confirmed the annotation by comparison with databases and literature (Lewis et al., 1998; Fossen et al., 2003; Ieri et al., 2011; Eichhorn et al., 2011; Navarre et al., 2013; López-Cobo et al., 2014), by using authentic standards for HCCs, and by HPLC-HRMS accurate mass for six compounds (Table 3). We propose the identification of two miscellaneous (sinapine, salicylic acid glucoside, one remains unknown), four polyamines (caffeoyl putrescine, feruloyl putrescine, bis dihydrocaffeoyl spermidine, tris dihydrocaffeoyl spermine), four HCCs (3-CGA, 4-CGA, 5-CGA, cis-5-CGA), ten anthocyanins (four peonidin derivatives, one cyanidin derivative, five pelargonidin derivatives, plus two pelargonidin hexoses without identification), two non-anthocyanin flavonoids (eriodictyol hexose, rutin, plus three putative flavonoids with phenolic acids), and two glycoalkaloids (solanine, chaconine). Moreover, we tentatively annotated isomers (identical mass spectra and different RT), which are the case of the four CGA isomers (3-CGA, 4-CGA, 5-CGA, cis-5-CGA) and the six pelargonidin derivatives indicated as Form 1 (F1) and Form 2 (F2) (Table 3).

**Table 3.**
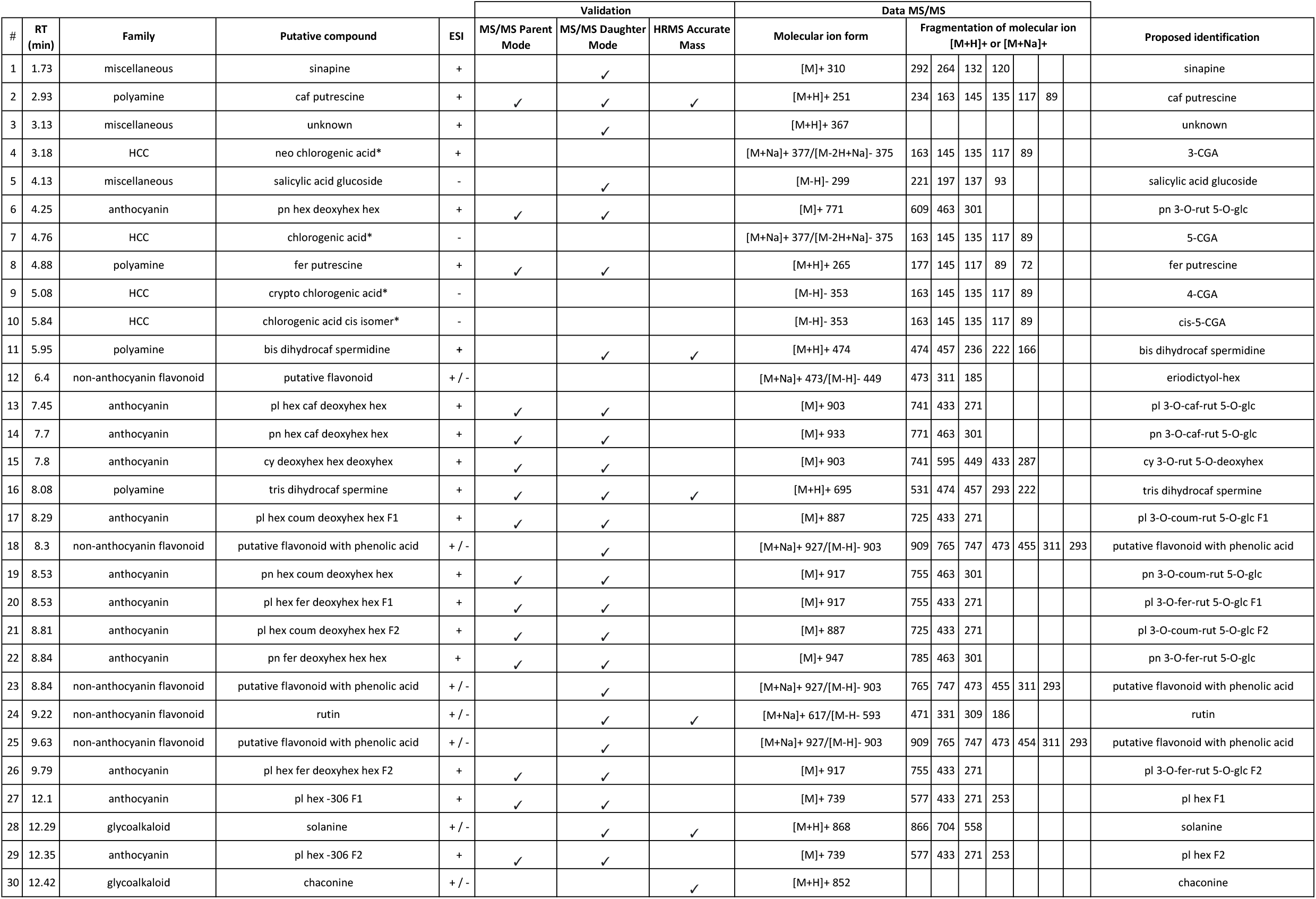
Identification data of the 30 compounds detected by HPLC-ESI-MS/MS in potato samples. #, peak number; *, validated by standard; caf, caffeoyl; coum, coumaroyl; cy, cyanidin; deoxyhex: deoxyhexose; ESI, electrospray ionization; fer, feruloyl; F1, form 1; F2, form 2; glc, glucose; hex, hexose; pl, pelargonidin; pn, peonidin; RT, retention time; rut, rutinoside.

### 3.4. Semi-quantitative HPLC-ESI-MS/MS analysis

Total ion current chromatograms (TICs) show that the profiles of TEs are similar to the profiles of AFs, except between RT 3 and 6 min, where TICs of TEs are similar to those of OFs (Supplemental Figures 4-6). Therefore, OFs were rich in the four HCCs, whereas AFs were rich in other compounds.

The semi-quantification results (μg apigenin equivalents/g DW) of the 30 compounds detected by mass spectrometry in TEs are shown in Table 4. Among families, we observed that HCCs, polyamines, and glycoalkaloids were found in higher concentrations than flavonoids in potato tubers. In general, HCCs presented higher concentrations in the skin than in the flesh (as was also found by HPLC-DAD). Among polyamines, caffeoyl and feruloyl putrescine levels were similar between skin and flesh. On the contrary, bis dihydrocaffeoyl spermidine was more abundant in the skin than in the flesh, and tri dihydrocaffeoyl spermine was only found in skin. Miscellaneous metabolites (sinapine, salicylic acid glucoside and unknown) were generally more abundant in flesh than skin. Among anthocyanins, peonidin and pelargonidin derivatives were found in both skin and flesh of Santa María, and in skin of Waicha and Moradita. Non-anthocyanin flavonoids were principally found in Santa María skin and flesh. Among glycoalkaloids, solanine and chaconine levels were more abundant in the skin than flesh, and absent in Waicha flesh (Table 4).

**Table 4.**
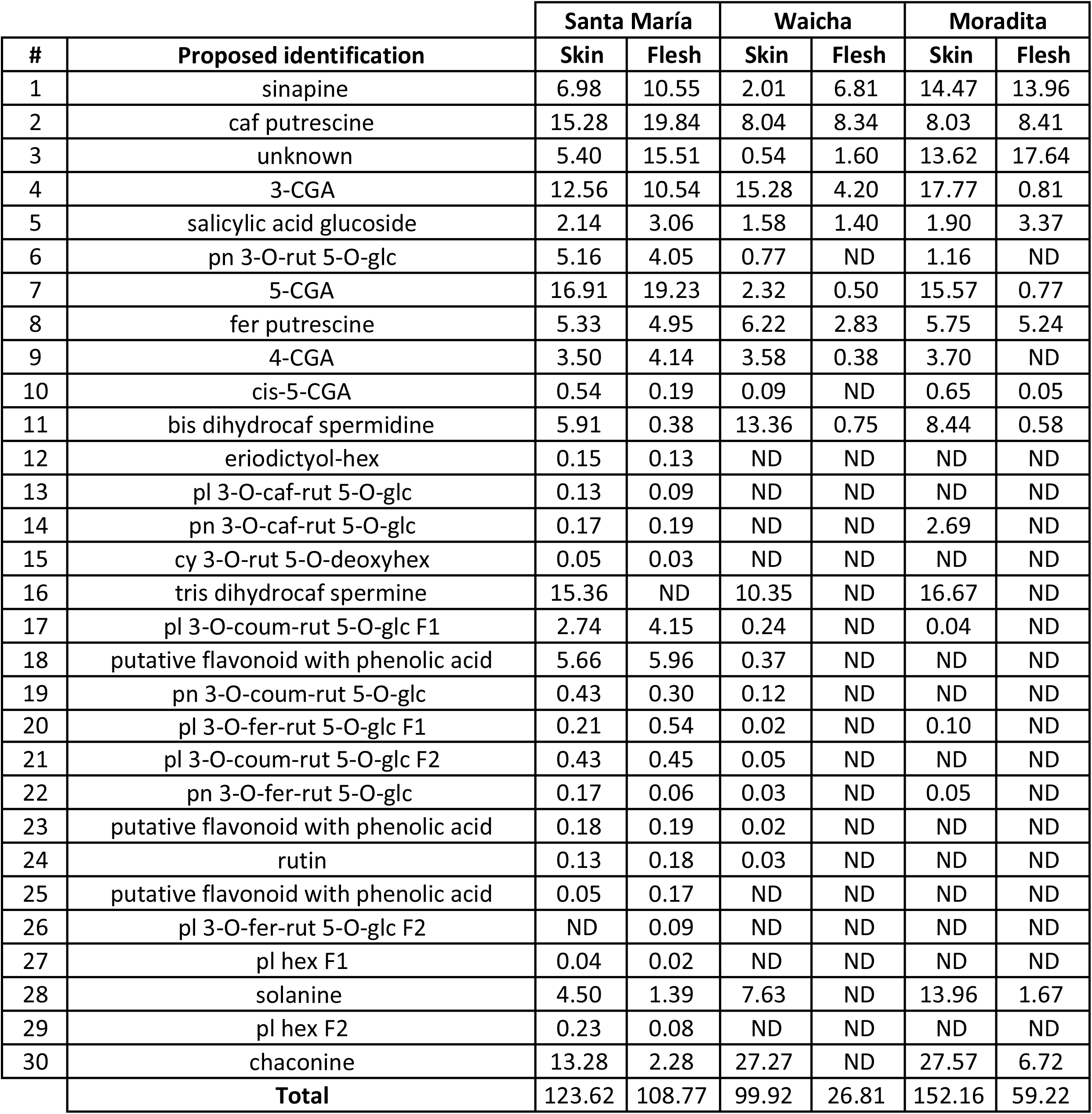
Content of individual compounds by HPLC-ESI-MS/MS in potato samples. Levels of metabolites were expressed as μg apigenin equivalents/g DW. ND, not detected.

| # | Proposed identification | Santa María |  | Waicha |  | Moradita |  |
| --- | --- | --- | --- | --- | --- | --- | --- |
|  |  | Skin | Flesh | Skin | Flesh | Skin | Flesh |
| 1 | sinapine | 6.98 | 10.55 | 2.01 | 6.81 | 14.47 | 13.96 |
| 2 | caf putrescine | 15.28 | 19.84 | 8.04 | 8.34 | 8.03 | 8.41 |
| 3 | unknown | 5.40 | 15.51 | 0.54 | 1.60 | 13.62 | 17.64 |
| 4 | 3-CGA | 12.56 | 10.54 | 15.28 | 4.20 | 17.77 | 0.81 |
| 5 | salicylic acid glucoside | 2.14 | 3.06 | 1.58 | 1.40 | 1.90 | 3.37 |
| 6 | pn 3-O-rut 5-O-glc | 5.16 | 4.05 | 0.77 | ND | 1.16 | ND |
| 7 | 5-CGA | 16.91 | 19.23 | 2.32 | 0.50 | 15.57 | 0.77 |
| 8 | fer putrescine | 5.33 | 4.95 | 6.22 | 2.83 | 5.75 | 5.24 |
| 9 | 4-CGA | 3.50 | 4.14 | 3.58 | 0.38 | 3.70 | ND |
| 10 | cis-5-CGA | 0.54 | 0.19 | 0.09 | ND | 0.65 | 0.05 |
| 11 | bis dihydrocaf spermidine | 5.91 | 0.38 | 13.36 | 0.75 | 8.44 | 0.58 |
| 12 | eriodictyol-hex | 0.15 | 0.13 | ND | ND | ND | ND |
| 13 | pl 3-O-caf-rut 5-O-glc | 0.13 | 0.09 | ND | ND | ND | ND |
| 14 | pn 3-O-caf-rut 5-O-glc | 0.17 | 0.19 | ND | ND | 2.69 | ND |
| 15 | cy 3-O-rut 5-O-deoxyhex | 0.05 | 0.03 | ND | ND | ND | ND |
| 16 | tris dihydrocaf spermine | 15.36 | ND | 10.35 | ND | 16.67 | ND |
| 17 | pl 3-O-coum-rut 5-O-glc F1 | 2.74 | 4.15 | 0.24 | ND | 0.04 | ND |
| 18 | putative flavonoid with phenolic acid | 5.66 | 5.96 | 0.37 | ND | ND | ND |
| 19 | pn 3-O-coum-rut 5-O-glc | 0.43 | 0.30 | 0.12 | ND | ND | ND |
| 20 | pl 3-O-fer-rut 5-O-glc F1 | 0.21 | 0.54 | 0.02 | ND | 0.10 | ND |
| 21 | pl 3-O-coum-rut 5-O-glc F2 | 0.43 | 0.45 | 0.05 | ND | ND | ND |
| 22 | pn 3-O-fer-rut 5-O-glc | 0.17 | 0.06 | 0.03 | ND | 0.05 | ND |
| 23 | putative flavonoid with phenolic acid | 0.18 | 0.19 | 0.02 | ND | ND | ND |
| 24 | rutin | 0.13 | 0.18 | 0.03 | ND | ND | ND |
| 25 | putative flavonoid with phenolic acid | 0.05 | 0.17 | ND | ND | ND | ND |
| 26 | pl 3-O-fer-rut 5-O-glc F2 | ND | 0.09 | ND | ND | ND | ND |
| 27 | pl hex F1 | 0.04 | 0.02 | ND | ND | ND | ND |
| 28 | solanine | 4.50 | 1.39 | 7.63 | ND | 13.96 | 1.67 |
| 29 | pl hex F2 | 0.23 | 0.08 | ND | ND | ND | ND |
| 30 | chaconine | 13.28 | 2.28 | 27.27 | ND | 27.57 | 6.72 |
| Total |  | 123.62 | 108.77 | 99.92 | 26.81 | 152.16 | 59.22 |

### 3.5. Total phenolics and selection of cytotoxic doses of TEs

To calculate the doses for the treatments in cells, we performed the Folin-Ciocalteu reaction with the different TEs. As expected, results showed that the skin extracts had more phenolic concentration than the flesh extracts. Among cultivars, Santa María total phenolic content is the highest, followed by Moradita and Waicha (Table 5).

**Table 5.** Total phenolics content in TE by the Folin-Ciocalteu assay. Results are expressed as μg 5-CGA equivalents/mL.

| Cultivar | Tissue | Total phenolics |
| --- | --- | --- |
| | | ($\mu\text{g}$ 5-CGA equivalents/mL) |
| Santa María | Skin | 16480 |
|  | Flesh | 6280 |
| Waicha | Skin | 3640 |
|  | Flesh | 990 |
| Moradita | Skin | 9160 |
|  | Flesh | 1500 |

We then used doses to an approximate 50% of cytotoxic activity in human neuroblastoma SH-SY5Y cells based on the MTT assays previously reported by our group (Silveyra et al., 2018). However, Figure 2 showed a minor effect on % cell viability than expected. Therefore, we decided to continue with 100 μg/mL Santa María skin, 400 μg/mL Santa María flesh, 10 μg/mL Waicha skin, 300 μg/mL Waicha flesh, 30 μg/mL Moradita skin, and 60 μg/mL Moradita flesh.

**Figure 2.**
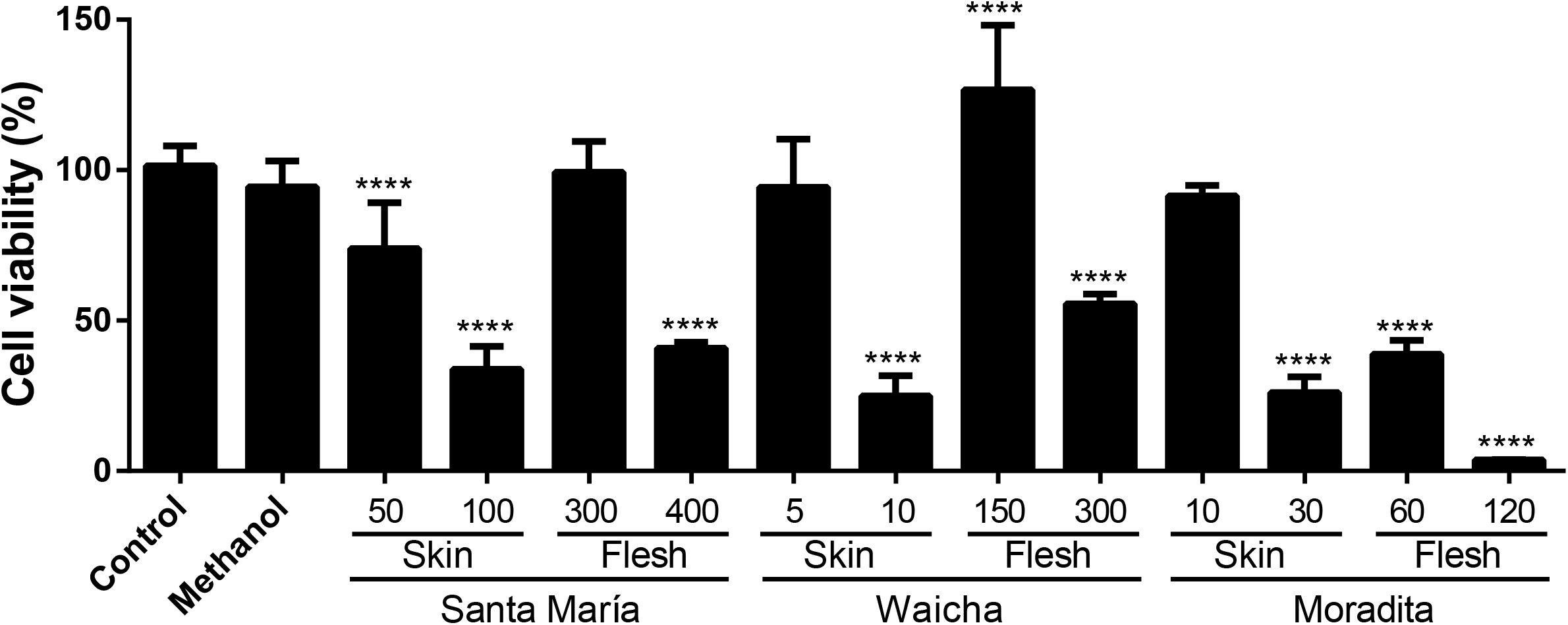
Cytotoxic activity of TEs from skin and flesh of Santa María, Waicha, and Moradita cultivars against human neuroblastoma SH-SY5Y cells. Doses are expressed as μg 5-CGA equivalents/mL. Values represent the average ± SD from at least three experiments with three technical repetitions. One way ANOVA multiple comparisons was followed by Dunnet’s test. ****, significant differences with P <0.0001; Control, non-treated cells; Methanol, cells treated with methanol ≤3% v/v.

### 3.6. Treatments of cells with TEs, OFs, and AFs: microscopy

We treated SH-SY5Y cells with the TEs, fractions, and the sum OF+AF. Before measuring cell viability by MTT assay, we observed cells under the microscope. Representative photographs of cells treated during 18 h with Waicha skin and flesh (10 and 300 μg/mL, respectively) are shown in Figure 3. Waicha was chosen to visualize the morphological changes because the skin extract exerted the highest cytotoxic activity with the lowest phenolic concentration (Figure 2).

**Figure 3.**
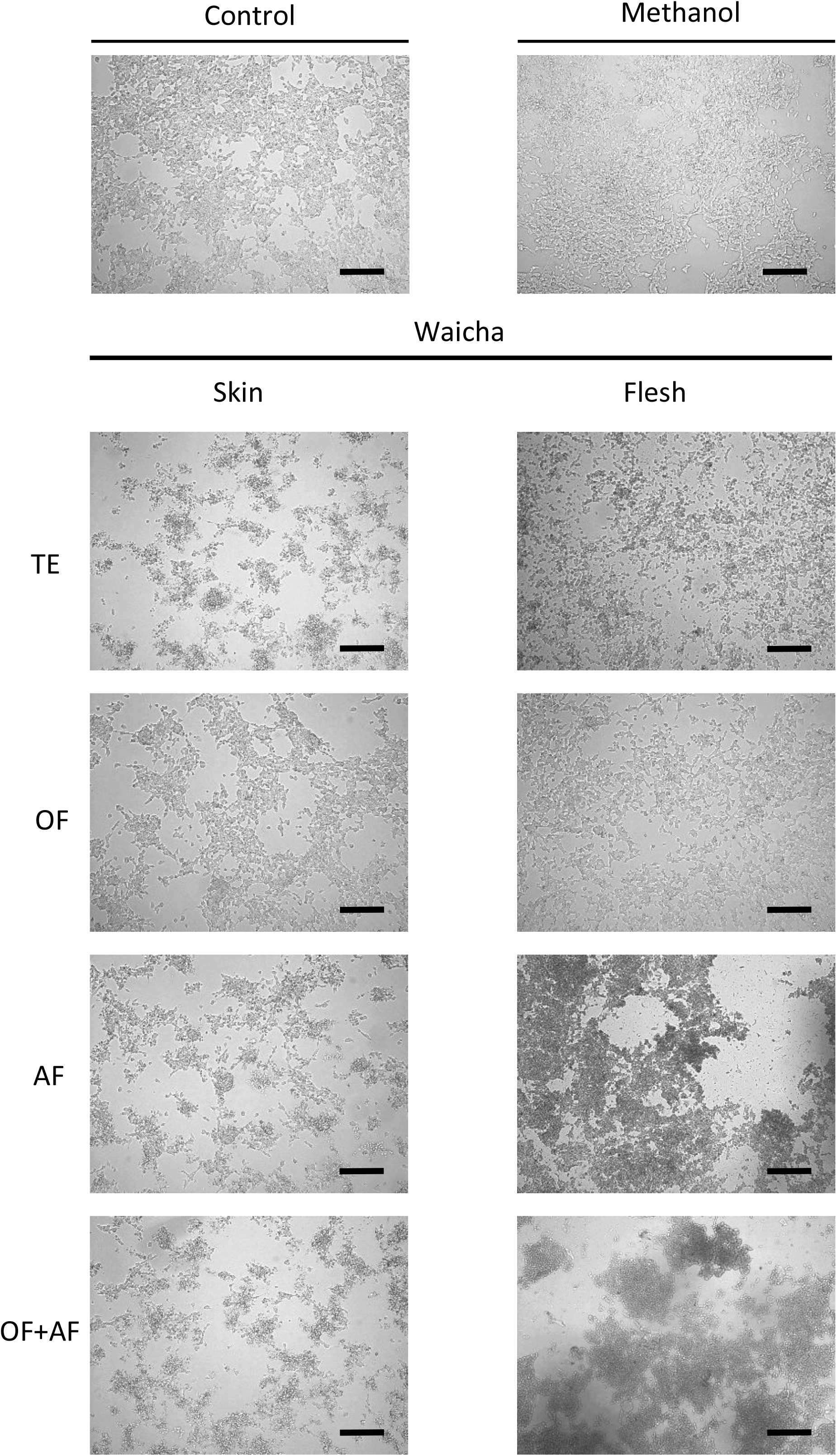
SH-SY5Y cells were treated for 18 h with TE, OF, AF, and OF+AF from Waicha skin (10 μg/mL) and flesh (300 μg/mL). Control, non-treated cells; Methanol, cells treated with methanol ≤3% v/v. Cells were examined under inverted microscopy (10X). The bar represents 200 μm.

The typical morphology of SH-SY5Y cells is neuronal, with a starry form. They grow adhered to the substrate and in a minor quantity in suspension. Both control and methanol treatments showed a pattern consistent with normal cell morphology (Figure 3). Cells treated with TEs from Waicha skin and flesh showed morphological changes compared to controls: they were reduced in size, had a rounded form, and were aggregated in cumulus. However, cells were similar to the controls in treatments with OFs from both tissues. Treatments with AFs exerted a similar effect to those with OF+AF, showing cell damage in both skin and flesh extracts (Figure 3).

### 3.7. Cytotoxic assays with TEs, OFs, and AFs: cell viability

After visualizing the cells, we quantified the cytotoxic effect by the MTT assay. All treatments with TEs resulted in a cell viability ≤50% (Figure 4). Treatment with OFs did not cause cell death and in Waicha provoked an increment in cellular metabolism compared to controls. AFs were cytotoxic in all cases, even superior to the corresponding TE in Santa María flesh. Except for Santa María flesh, treatments with OF+AF showed diminished cell viability, with values similar to the respective TEs (Figure 4). Values obtained for Waicha skin and flesh are consistent with the phenotypes observed in Figure 3.

**Figure 4.**
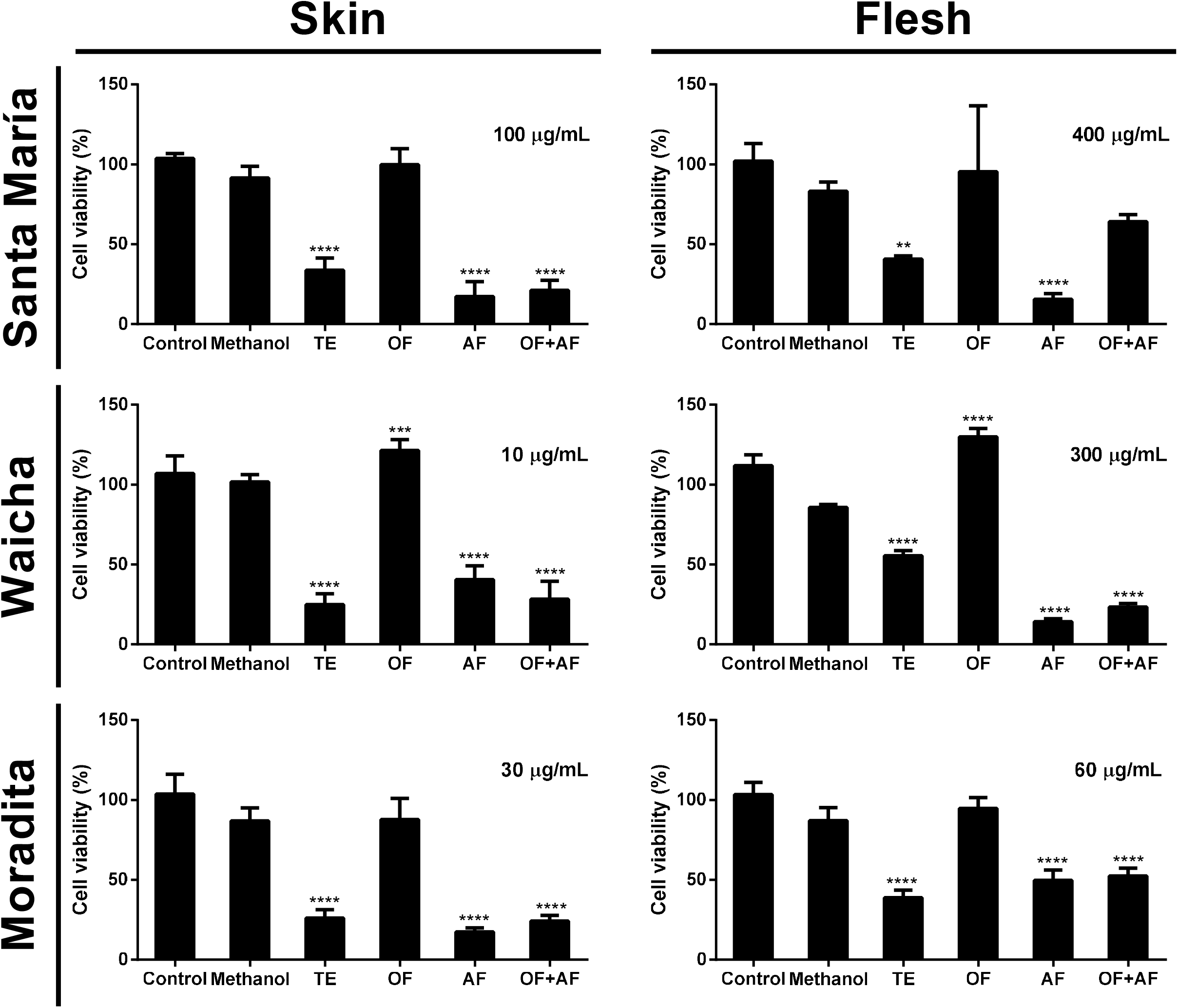
Cytotoxic activity of TE, OF, AF, and OF+AF from skin and flesh of Santa María, Waicha, and Moradita cultivars against human neuroblastoma SH-SY5Y cells. Doses are expressed as μg 5-CGA equivalents/mL. Values represent the average ± SD from at least three experiments with three technical repetitions. One way ANOVA multiple comparisons was followed by Dunnet’s test. **, significant differences with P<0.005; ***, significant differences with P<0.001; ****, significant differences with P<0.0001; Control, non-treated cells; Methanol, cells treated with methanol ≤3% v/v.

### 3.8. Correlation analysis between the content of metabolites and cell viability

In order to reveal the contribution of compounds to the cytotoxic activity, we analyzed the Pearson correlation coefficients between the mean values of metabolites levels and of % cell viability (Figure 5). We were particularly interested in the compounds present in the AFs. Notably, three polyamines (feruloyl putrescine, bis dihydrocaffeoyl spermidine, and tris dihydrocaffeoyl spermine) (Figure 5A) and two glycoalkaloids (solanine and chaconine) (Figure 5B) exerted negative and significant correlation (R^2^≥0.5743). The correlation was more evident for feruloyl putrescine (R^2^=0.9265) and chaconine (R^2^=0.8334). In addition, the sum of the content of the three polyamines and the two glycoalkaloids negatively correlated with cell viability (R^2^=0.8122, Figure 5C). Results indicate that a higher content of those five metabolites would be accompanied by lesser cell viability, or in other words, by a higher cytotoxic activity. Correlation analysis for the other compounds in AFs was not significant (data not shown).

**Figure 5.**
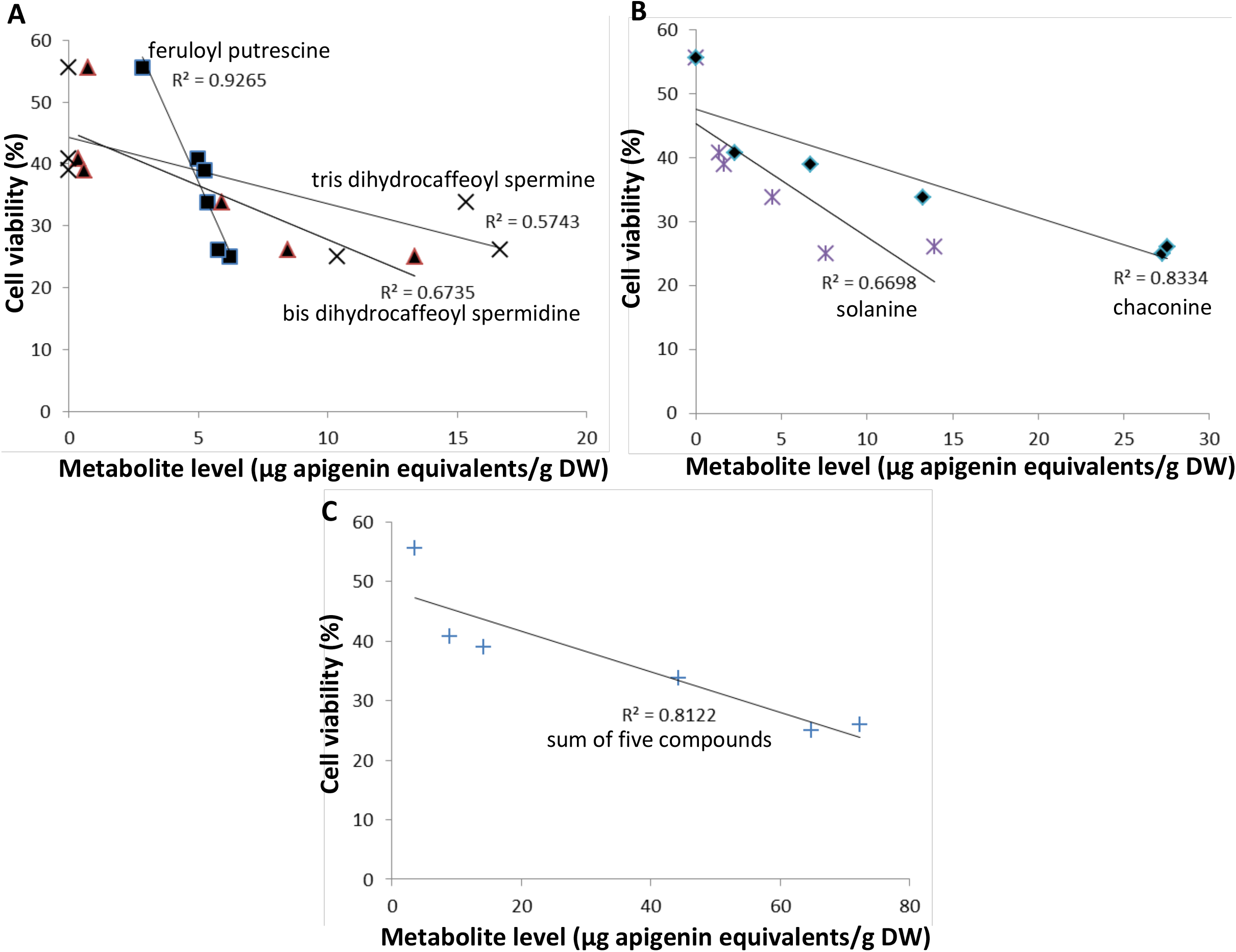
Correlation analysis between the content of individual metabolites and % cell viability in human neuroblastoma SH-SY5Y cells. A, Pearson correlation coefficients for the polyamines feruloyl putrescine, bis dihydrocaffeoyl spermidine, and tris dihydrocaffeoyl spermine. B, Pearson correlation coefficients for the glycoalkaloids solanine and chaconine. C, Pearson 697 correlation coefficients for the sum of the three polyamines and the two 698 glycoalkaloids.

## 4. Discussion

This work deepens into the cytotoxic activity of phenolic extracts from Andean potatoes via liquid-liquid fractionation in order to identify the compounds responsible for that activity.

TEs of skin and flesh of three Andean potato cultivars were successfully fractionated into OFs and AFs. Results of HPLC-DAD (and HPLC-ESI-MS/MS) corroborated the expected composition of both OFs and AFs. Fractionation was good but not optimal (considering the remaining compounds in each fraction), being AFs purer than OFs. This result indicates that more incubation time and/or more repetitions of successive extractions with ethyl acetate could be required for a better fractionation. Further, the differences between the sum of compounds in OF+AF compared to TE values could be attributed to slight differences in resuspension volumes of the samples after complete dryness, and/or to errors during HPLC-DAD measurement (coming from the equipment or from dilutions performed to avoid column saturation).

HPLC-DAD results indicate that the main metabolite in all potato cultivars and tissues is 5-CGA, with a significant presence of CA in pigmented tissues. This result is consistent with previous reports that indicate that 5-CGA is the most abundant phenolic acid in potatoes, followed by CA (Navarre et al., 2011; Deusser et al., 2012). HPLC-ESI-MS/MS results indicate that the main metabolites are 5-CGA, 3-CGA (accurate mass and RT confirmed with standards), and caffeoyl putrescine; whereas tris dihydrocaffeoyl spermine and chaconine levels are also significantly in skin.

Agreeing with the literature (Bontempo et al., 2013; Valiñas et al., 2017), our analysis revealed that red tuber tissues (such as Santa María skin and flesh) contain mainly pelargonidin and peonidin derivatives, whereas purple tuber tissues (such as Moradita skin) harbors mostly derivatives of petunidin, malvidin (both found only by HPLC-DAD) and peonidin. No anthocyanidins were detected in Waicha and Moradita fleshes, which is expected since they lack pigmentation. The discrepancy between anthocyanidin/anthocyanin compositions by HPLC-DAD/HPLC-ESI-MS/MS could be due to differences in sample treatments (resuspension solvent) and/or chromatographic conditions (column, mobile phases). Some differences could be also attributed to the acid hydrolysis prior to HPLC-DAD since all aglycones are recognized as only one peak (e.g. pelargonidin). In contrast, no acid hydrolysis is needed for mass spectrometry analysis because each peak is semi-quantitated (HPLC-ESI-MS/MS) and individually identified (HPLC-HRMS).

HPLC-DAD chromatograms showed that besides the quantified compounds, there are some unidentified peaks. HPLC-DAD technique is limited due to the lack of authentic standards for compound identification by comparison with absorption spectra and RT. By contrast, HPLC-ESI-MS/MS and HPLC-HRMS offer a more complete and accurate identification of metabolites. It should be noted that the levels of specific metabolites (e.g. CGA isomers) by HPLC-ESI-MS/MS are lesser than the ones obtained by HPLC-DAD. However, cis-5-CGA could only be quantified by mass spectrometry. Notably, metabolomic profiles by mass spectrometry have not been described for any of the genotypes included in our study. Also, few comparisons have been made between potato tuber skin and flesh (López-Cobo et al., 2014; Oertel et al., 2017; Valiñas et al., 2017; Nogawa et al., 2019). To conclude, in our investigations, both techniques are complementary and necessary.

HPLC-DAD analysis underestimates the content of total phenolics because it only quantifies some of the present compounds. Folin-Ciocalteu assay overestimates the content of total phenolics because it reacts with compounds like vitamins, aminoacids, and thiol derivatives (Everette et al., 2010). Nevertheless, we found a significant positive Pearson correlation between HPLC-DAD and Folin-Ciocalteu data (R^2^=0.8612).

The assays for selecting cytotoxic doses in human neuroblastoma SH-SY5Y cells show a lesser activity of the extracts than expected, taking into account previous studies from our laboratory (Silveyra et al., 2018). This result could be due to the use of new stock of cells and/or changes in the reagents used for cell culture.

Together, results obtained by Folin-Ciocalteu and the assay of cytotoxic doses in SH-SY5Y cells, it can be concluded that a higher concentration of total phenolics does not imply a higher cytotoxic activity. For example, among the skins of the three cultivars, Waicha has the lesser total phenolics content (3640 μg/mL) and requires the lesser concentration (10 μg/mL) to induce cell mortality. Santa María skin has the highest level of total phenolics (16480 μg/mL) but requires the highest dose (100 μg/mL) to exert approximate 50% cell viability. A similar result was obtained for inhibiting cell proliferation in human breast cancer by potato extracts (Leo et al., 2008). Therefore, levels of specific compounds and/or interactions between different phytochemicals could be essential in inducing cytotoxic activity.

Concerning assays with SH-SY5Y cells, we observed a similar pattern for the TEs and their fractions in all the tested cultivars and tissues. OFs, rich in HCCs, were not cytotoxic in any case. In Waicha skin and flesh, they even provoked an increase in mitochondrial activity. Since treatments with 200-400 μg/mL of commercial 5-CGA were cytotoxic (Silveyra et al., 2018), we speculate that the absence of activity in OFs could be due to lesser doses of 5-CGA or to other components that antagonize the effect of 5-CGA. Conversely, AFs were cytotoxic in all cases, even in Waicha and Moradita fleshes which lack anthocyanins. Although numerous studies support the cytotoxic activity of potato extracts rich in anthocyanins against cancer cells (Reddivari et al., 2007; Bontempo et al., 2013; Vizzotto et al., 2014; Ombra et al., 2015), our study indicates that other components could be involved in inducing cell death. Results showed that treatments with OF+AF were cytotoxic with values near but not identical to the corresponding TEs. This result could be due to experimental errors generated by variations in the concentration when OFs and AFs are combined instead of TEs.

We used an untargeted mass spectrometry-metabolomics approach in order to gain information about bioactive compounds. Based on our observations, we can hypothesize that a combination of three polyamines (feruloyl putrescine, bis dihydrocaffeoyl spermidine and tris dihydrocaffeoyl spermine) and two glycoalkaloids (solanine and chaconine) are mainly responsible for the observed cytotoxic activity. This result reveals the nutraceutical properties and interactions of metabolites in a complex food matrix, like the potato. Since Santa María AF had major cytotoxic activity than the corresponding TE, we could even suggest that other compounds antagonize their effects. The next step could involve solid-phase fractionation to specifically recover glycoalkaloids, as was described by Sánchez Maldonado et al. 2014. Chaconine and solanine commercial standards induced 50% cytotoxicity against five mammalian cell lines at 2.9 μM and 18 μM, respectively (Nogawa et al., 2019). In addition, polyamines conjugated with phenolic acids (also called phenolamides) have been shown to exert anticancer properties in various cellular models (Roumani et al., 2020). In sum, future experiments should be directed to study the response of SH-SY5Y cells to treatments with different doses of individual or subgroups of bioactive metabolites. It would be interesting to determine individual compounds with cytotoxic activity by isolation with preparative HPLC, and the mechanisms of antitumoral activity. The cytotoxic effects of potato extracts involved the induction of apoptosis by pathways dependent and independent of caspases in prostate cancer cell lines (Reddivari et al., 2007). Other researchers have observed that extracts of Vitelotte cultivar cause dose-dependent inhibition of proliferation and induction of apoptosis in breast, cervix, prostate, and leukemia cancers (Bontempo et al., 2013).

This work contributes to potato revalorization as a functional food. It enlarges the knowledge of potato polyphenols’ bioactivity as promising sources of natural products with an added value for the food industry and human health.

## 5. Conclusions

The present study demonstrates that a complex interplay of polyphenolic compounds contributes to potato benefits. It also suggests that conjugated polyamines and glycoalkaloids are critical in inducing cytotoxicity against human neuroblastoma cells *in vitro*. Our results are of fundamental interest due to the nutritional, biological, and industrial contribution. Further studies are needed to identify and isolate the active compounds, determine their bioavailability, and confirm their neuroprotection *in vivo*.

## Contributions

M.L.L. and M.X.S. designed and instructed the research work. M.L.L., M.X.S., M.M.M., S.B., D-D.S-G. and F.P. performed the experiments. M.L.L. wrote the manuscript. M.L.L., M.X.S., and A.B.A. provided funding for this work. All authors commented on the manuscript and approved the final article.

## Acknowledgments

We thank M.S. Andrea Clausen, M.S. Ariana Digilio, and Lic. Patricia Suárez (Instituto Nacional de Tecnología Agropecuaria, Argentina) for assistance in obtaining the Andean tubers. We thank Lic. Daniela Villamonte (Instituto de Investigaciones Biológicas, Facultad de Ciencias Exactas y Naturales, Universidad Nacional de Mar del Plata, CONICET) for technical assistance in HPLC-DAD measurements. We are grateful to Drs. Tomás Falzone (Instituto de Biología Celular y Neurociencia “Prof. E. de Robertis”, Argentina) and Elena Avale (Instituto de Investigaciones en Ingeniería Genética y Biología Molecular “Dr. Héctor N. Torres”, Argentina) for providing the SH-SY5Y cell line. We thank Dr. Fernando Villarreal (Instituto de Investigaciones Biológicas, Facultad de Ciencias Exactas y Naturales, Universidad Nacional de Mar del Plata, CONICET) for suggesting substantial improvements to the manuscript.

This work was financially supported by grants from Agencia Nacional de Promoción Científica y Tecnológica (PICT 2018 3815, PICT 2018 0221), Consejo Nacional de Investigaciones Científicas y Técnicas (CONICET) (PIP 2015 0762) and Universidad Nacional de Mar del Plata (UNMdP, EXA 921/19). This work has benefited from the support of IJPB’s Plant Observatory technological platforms. The IJPB is financially supported by Saclay Plant Sciences (ANR-17-EUR-0007).

M.L.L., M.X.S., and A.B.A. are career members from CONICET. M.M.M. was an advanced undergraduate student, directed by M.L.L and codirected by M.X.S.

## Supporting information description

**Supplemental Figure 1**. HPLC-DAD chromatograms of TE, OF, and AF from Santa María skin and flesh. Profiles for HCCs were obtained at 320 nm and at 510 nm for anthocyanidins. 5-CGA, chlorogenic acid; CA, caffeic acid; FA, ferulic acid; Cy, cyanidin; Pl, pelargonidin; Pn, peonidin; Mv, malvidin.

**Supplemental Figure 2**. HPLC-DAD chromatograms of TE, OF, and AF from Waicha skin and flesh. Profiles for HCCs were obtained at 320 nm and at 510 nm for anthocyanidins. 5-CGA, chlorogenic acid; CA, caffeic acid; CouA, coumaric acid; FA, ferulic acid; Dp, delphinidin; Cy, cyanidin; Pt, petunidin; Pl, pelargonidin; Pn, peonidin; Mv, malvidin.

**Supplemental Figure 3**. HPLC-DAD chromatograms of TE, OF, and AF from Moradita skin and flesh. Profiles for HCCs were obtained at 320 nm and at 510 nm for anthocyanidins. 5-CGA, chlorogenic acid; CA, caffeic acid; CouA, coumaric acid; FA, ferulic acid; Dp, delphinidin; Cy, cyanidin; Pt, petunidin; Pl, pelargonidin; Pn, peonidin; Mv, malvidin.

**Supplemental Figure 4**. TIC chromatograms of TE, OF, and AF from Santa María skin and flesh. HPLC-ESI-MS/MS in positive ion mode.

**Supplemental Figure 5**. TIC chromatograms of TE, OF, and AF from Waicha skin and flesh. HPLC-ESI-MS/MS in positive ion mode.

**Supplemental Figure 6**. TIC chromatograms of TE, OF, and AF from Moradita skin and flesh. HPLC-ESI-MS/MS in positive ion mode.

