## Supplementary figures and images for "Metabolite profiling and cytotoxic activity of Andean potatoes: polyamines and glycoalkaloids as potential anticancer agents in human neuroblastoma cells *in vitro*"

### Supplemental Figures

#### Santa María skin

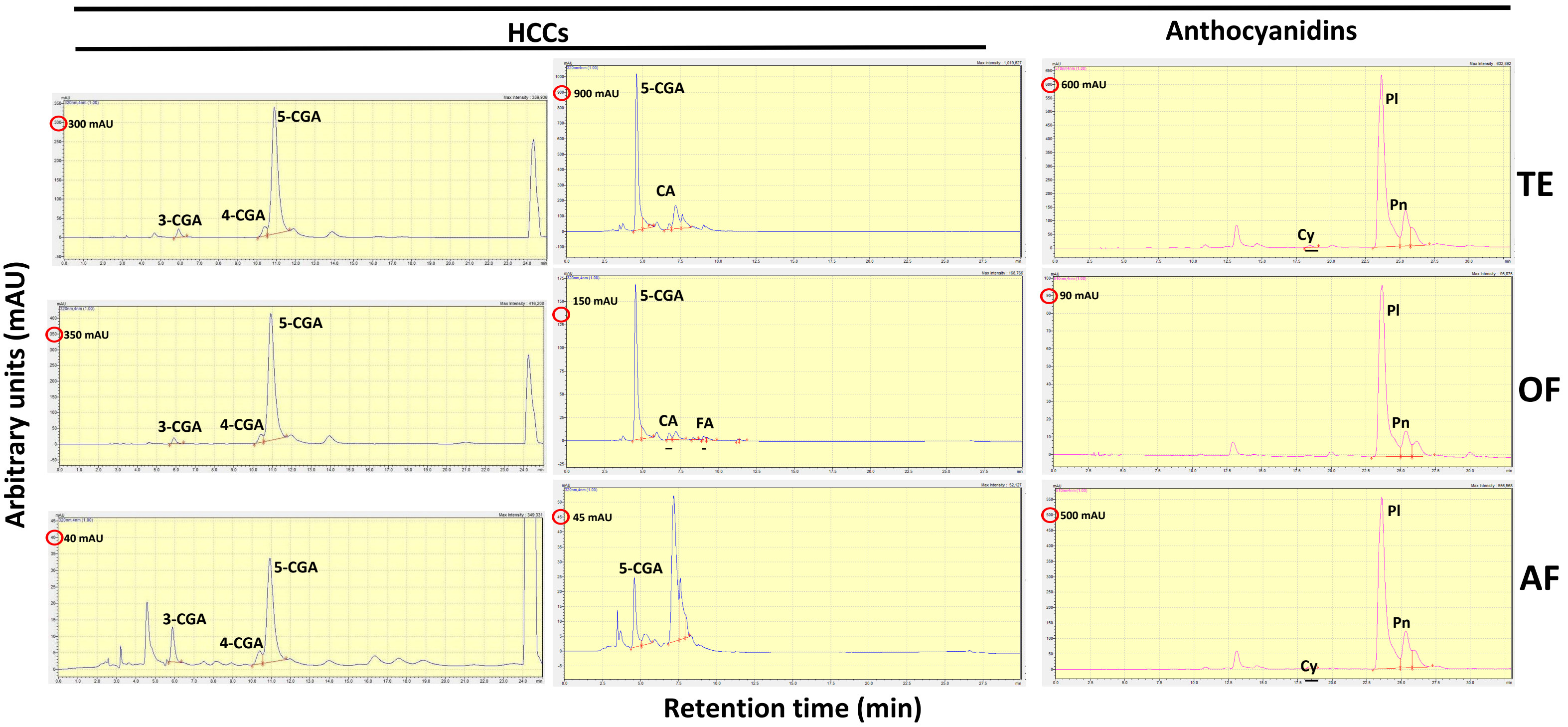


#### Santa María flesh

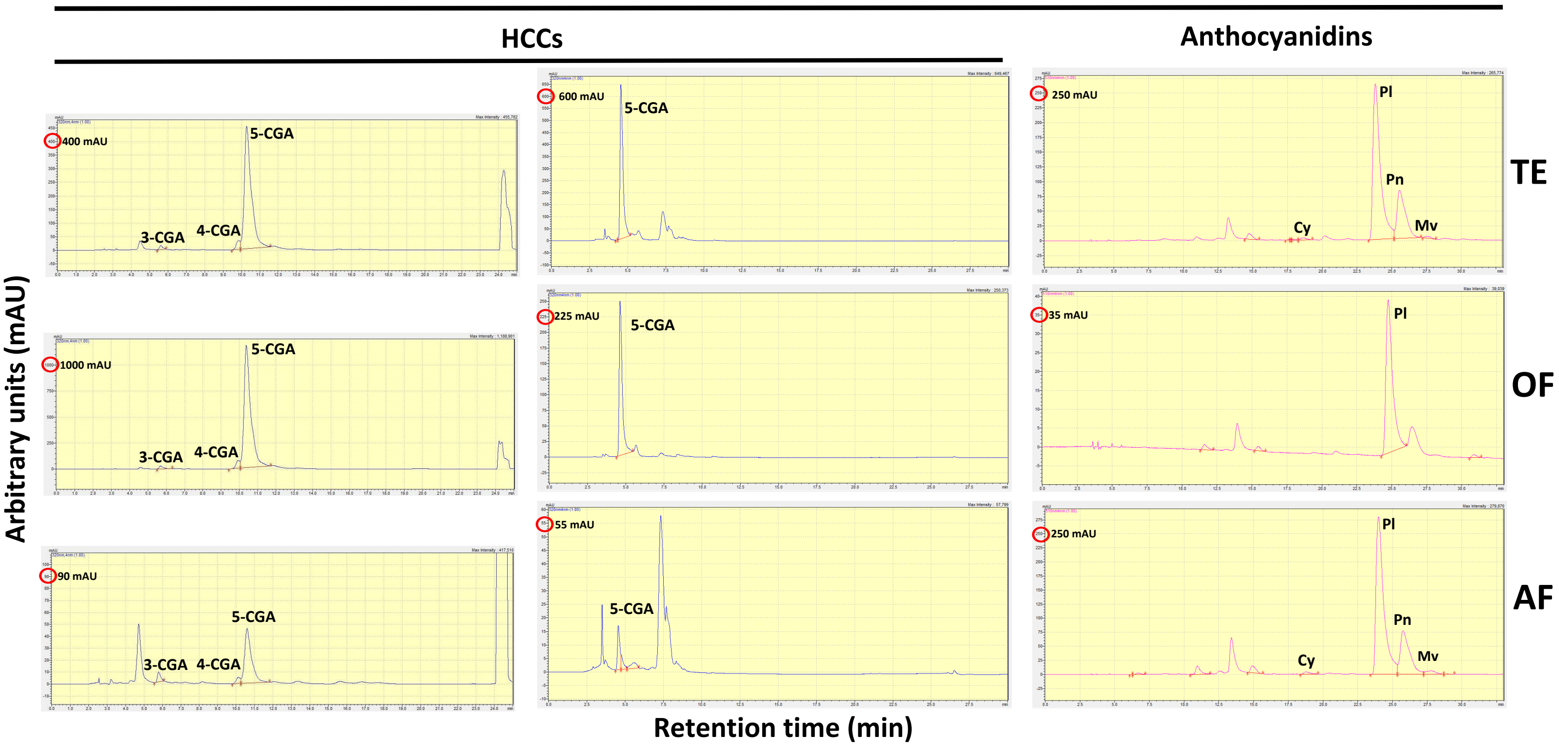

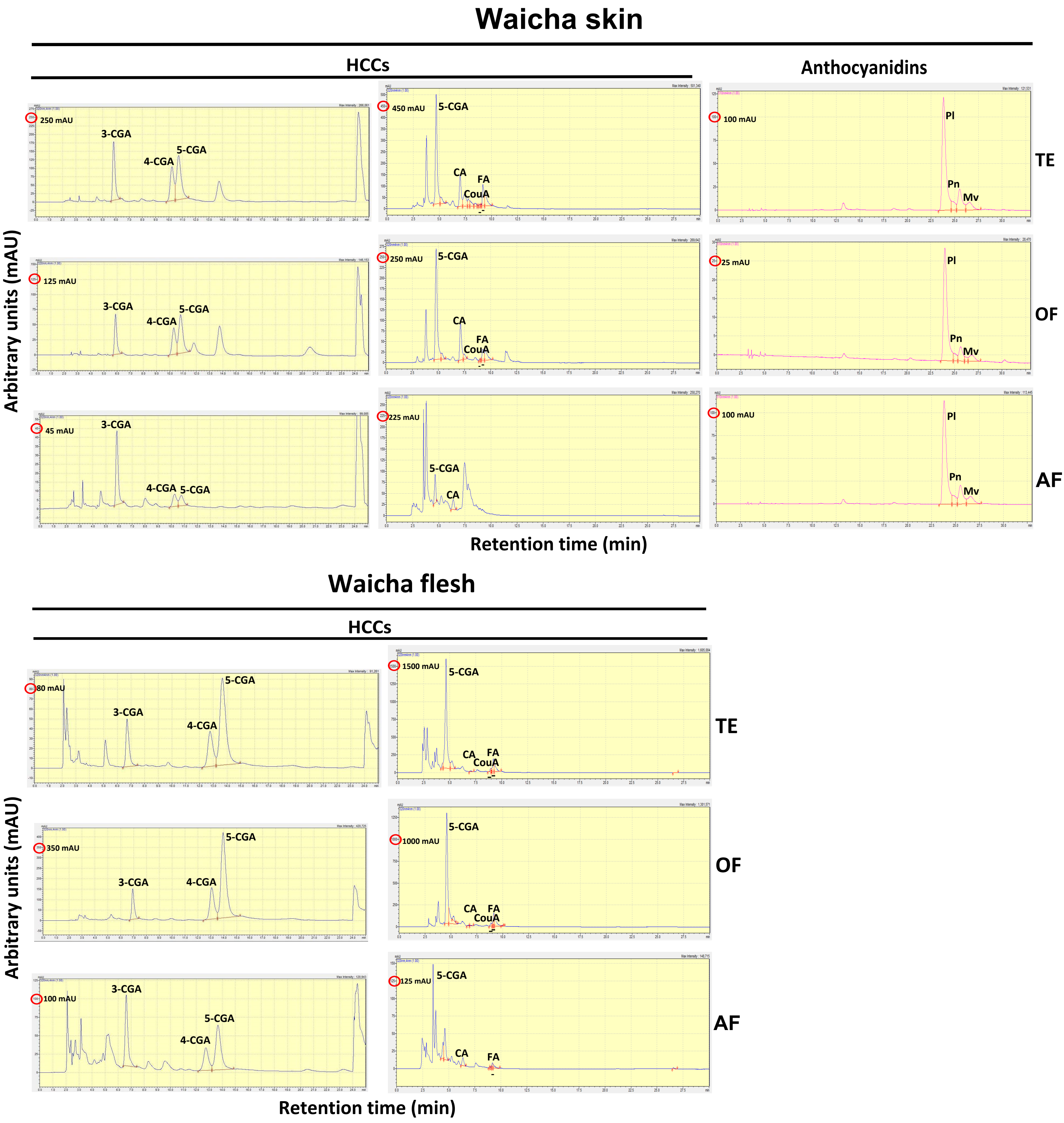

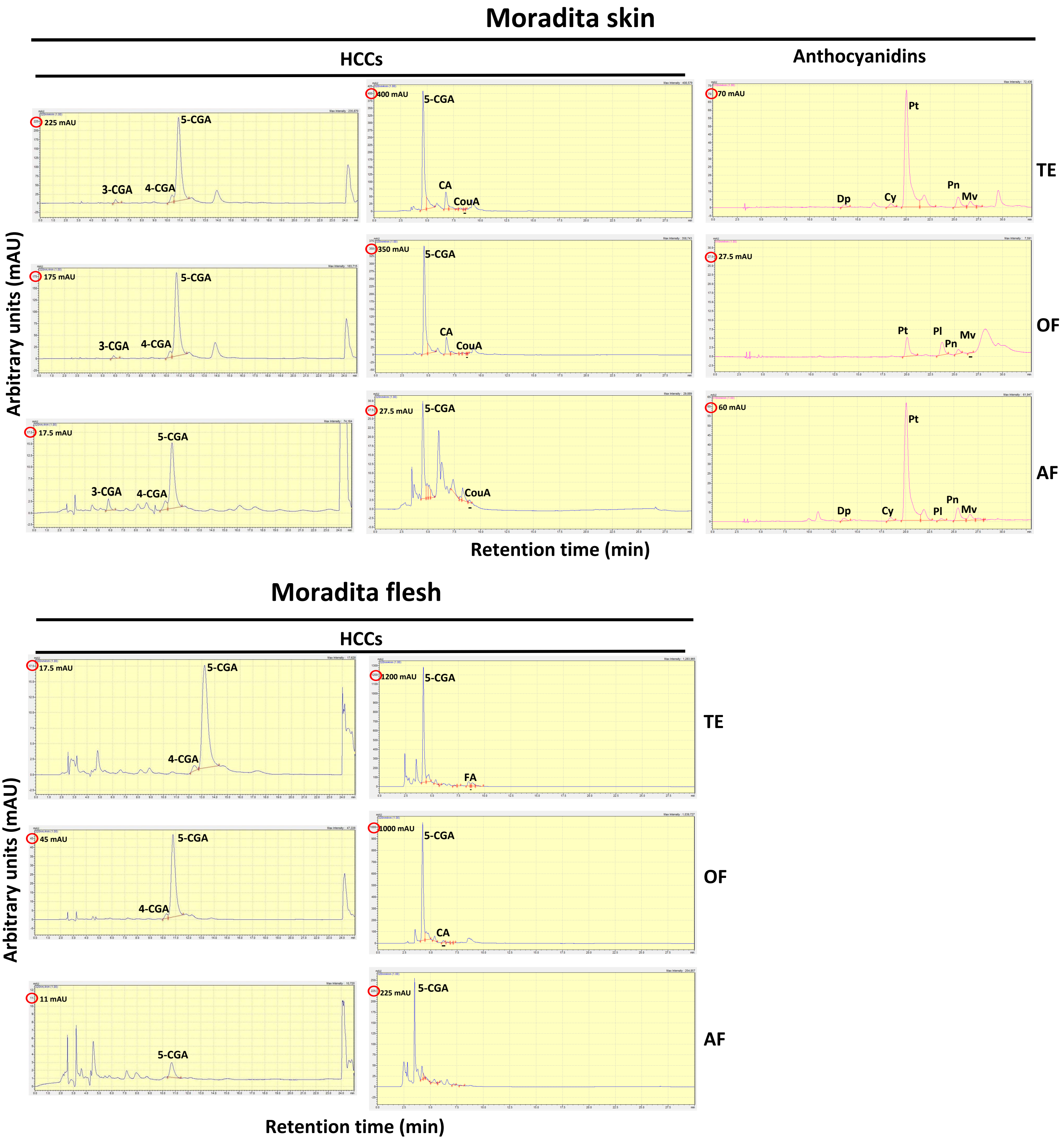

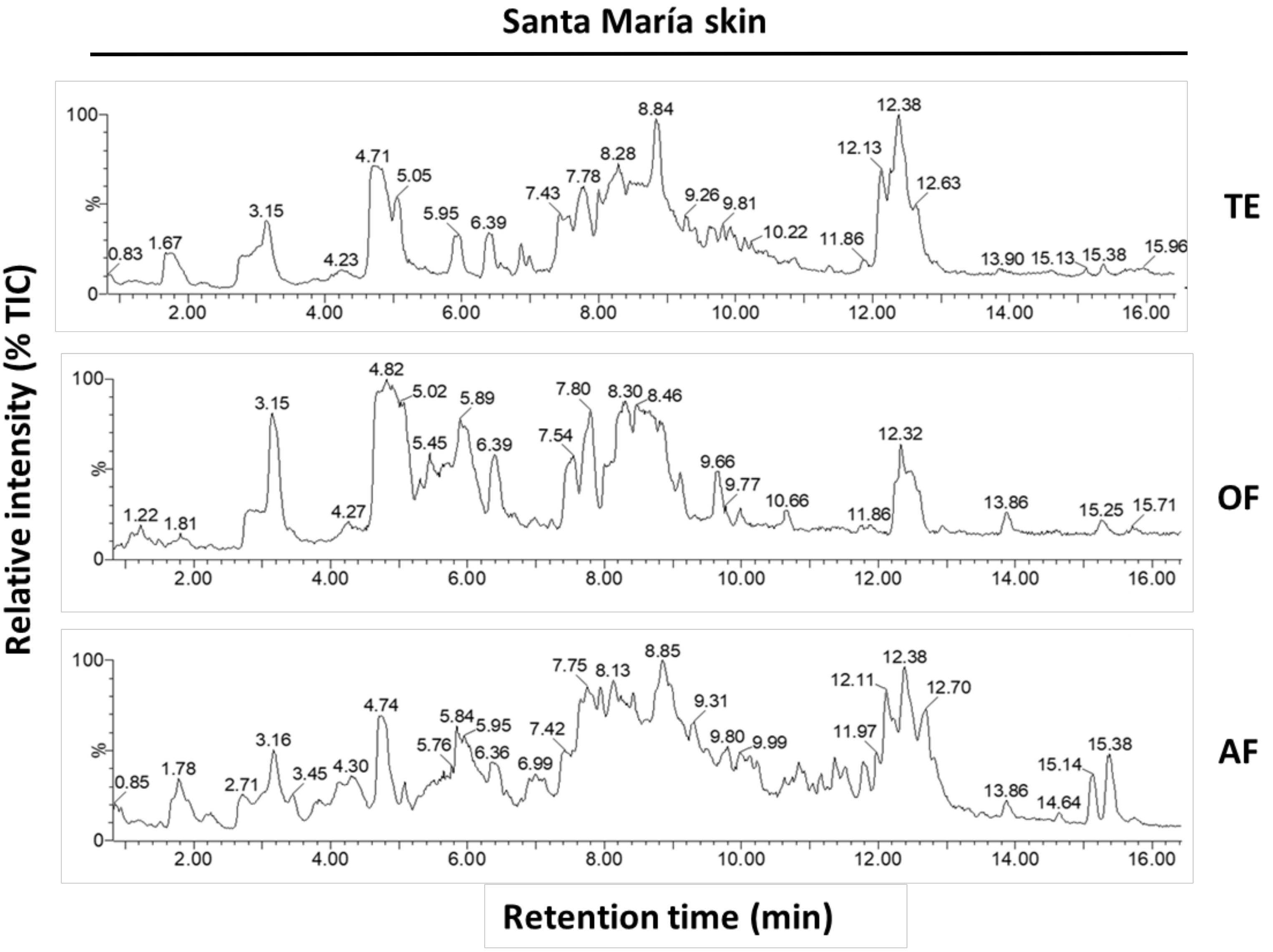

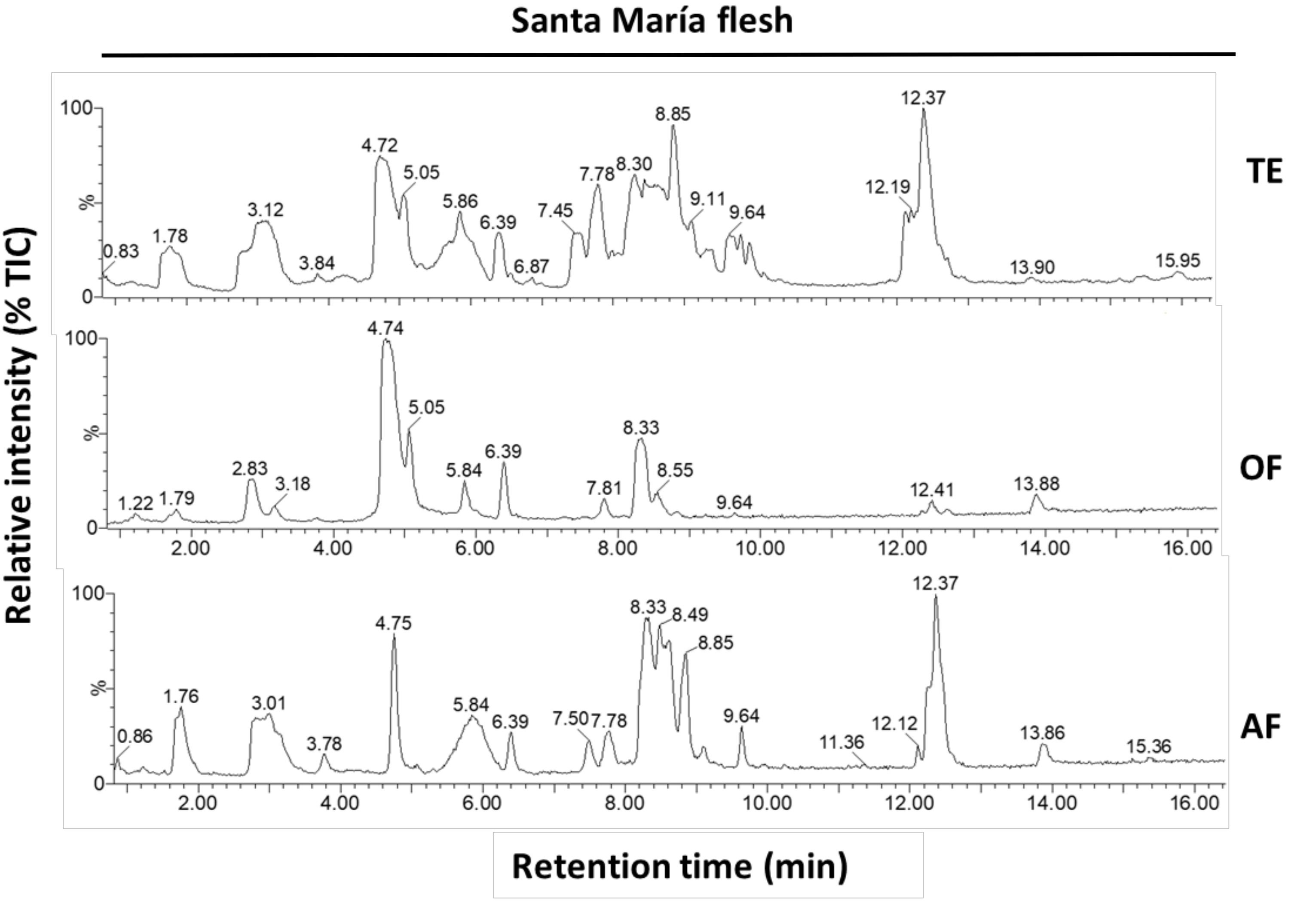

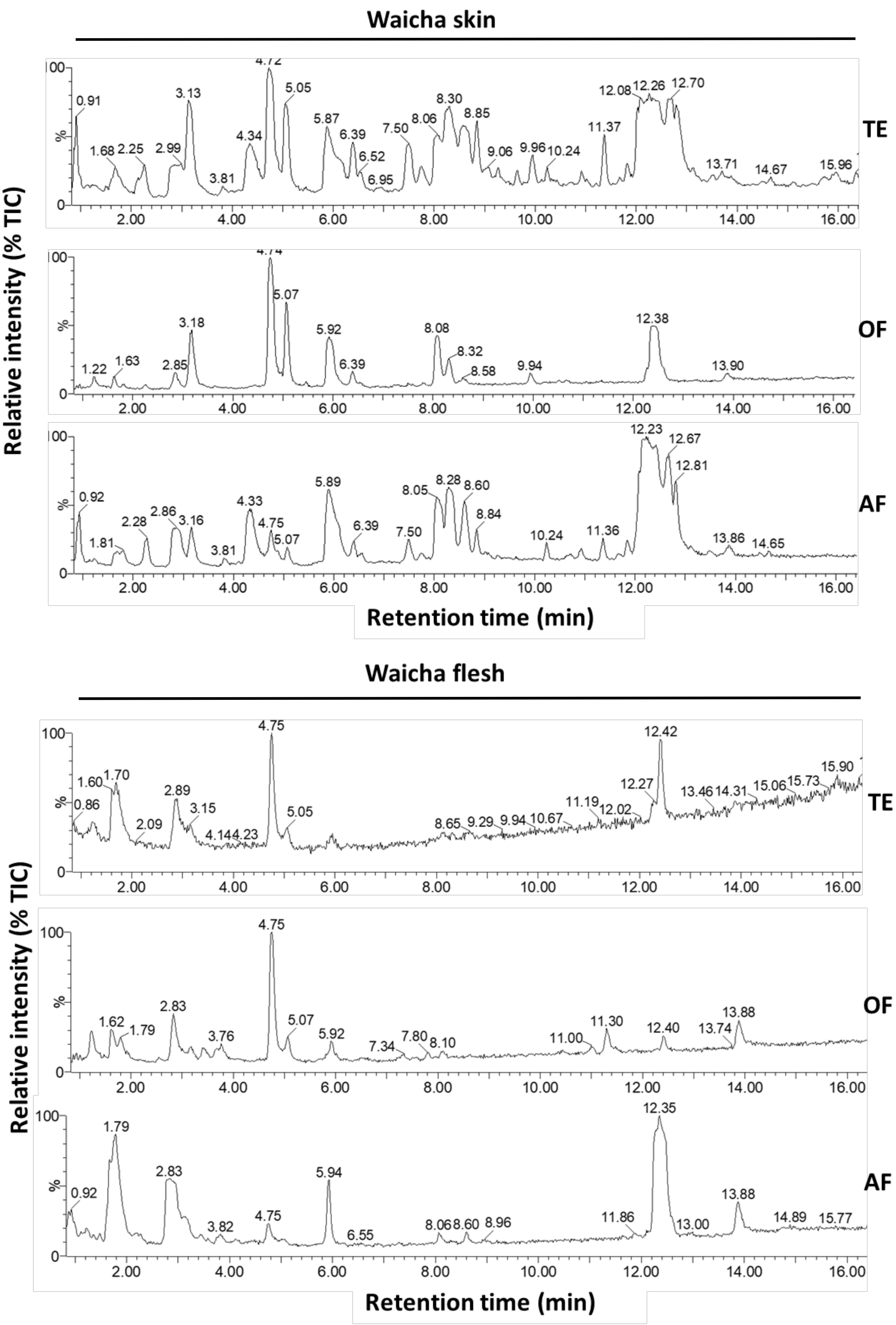

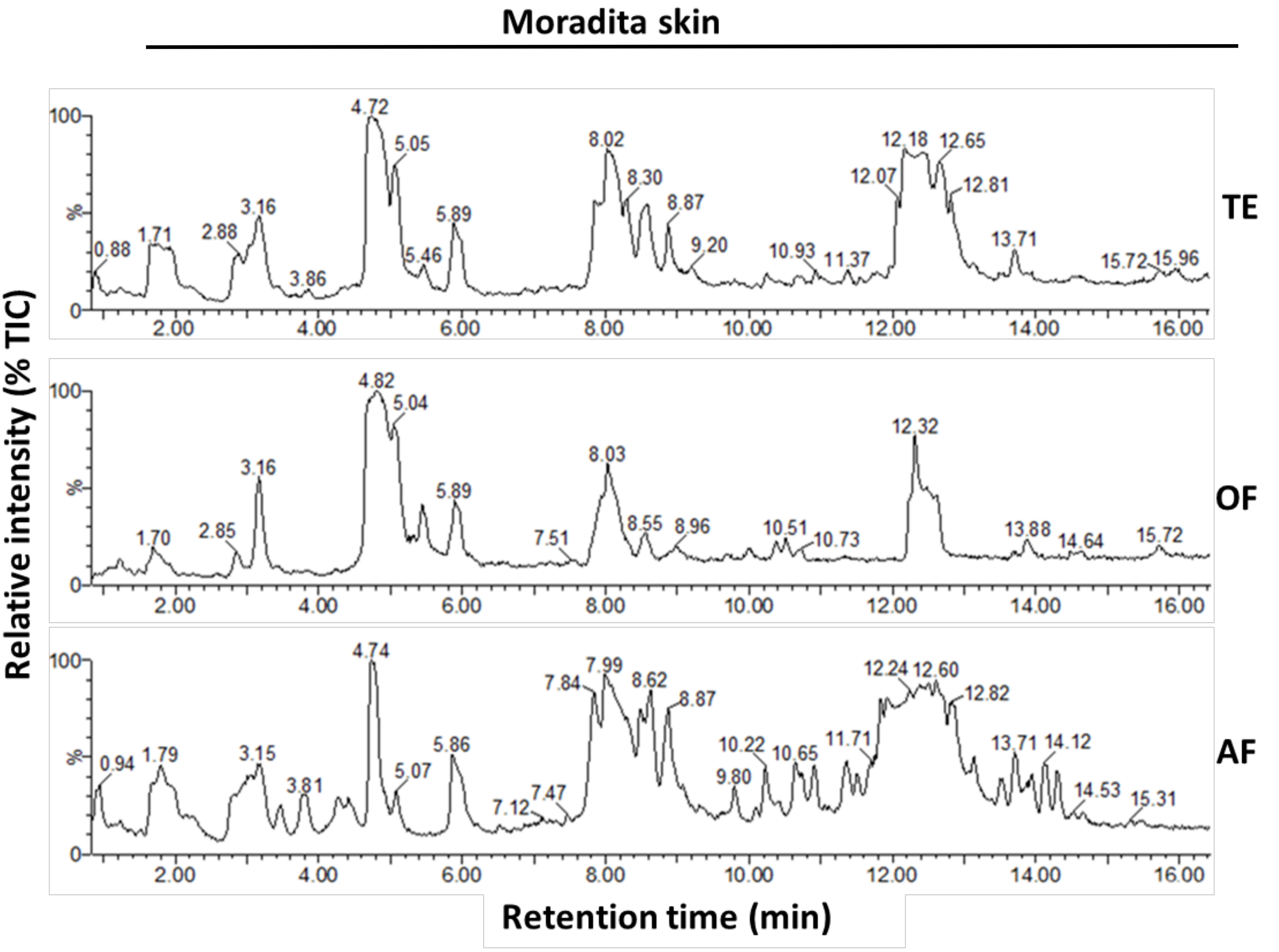

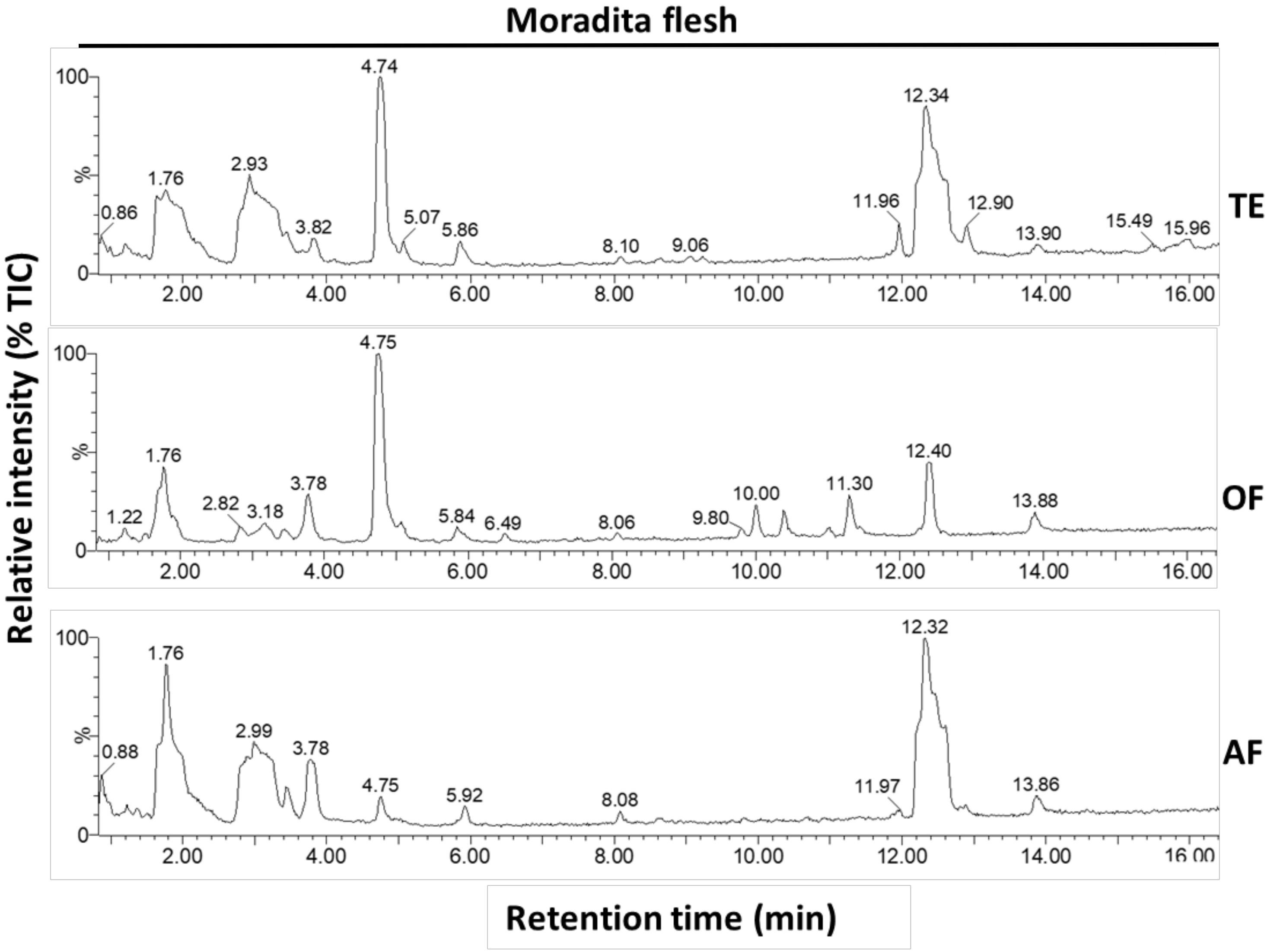
